# High-fat diet feeding reduces the expression of Rab3a, Rab3gap1, and Rab3Gap2 genes that are pivotal to neuronal exocytosis

**DOI:** 10.1101/2021.09.10.459820

**Authors:** Luana Assis Ferreira, Fernando Victor Martins Rubatino, Mariana Lacerda de Freitas, Leonardo Rossi de Oliveira, Célio Jose de Castro Junior, Fernanda Sarquis Jehee, Adriana Aparecida Bosco, Karla Silva Fernandes

**Affiliations:** Programa de Pós-graduação em Medicina e Biomedicina, Santa Casa de Misericórdia de Belo Horizonte – Ensino e Pesquisa, Belo Horizonte, MG, Brazil

**Keywords:** Rab3A, Rab3gap1, Rab3gap2, satiety, obesity, high-fat diet C57Bl/6 model, vesicle trafficking, neuronal exocytosis, arcuate nucleus, hypothalamus

## Abstract

The Rab3A and Rab3gaps are essential to the Ca^+2^-dependent neuronal exocytosis in the hypothalamus. The arcuate nucleus of the hypothalamus (ARC) controls food intake and energy expenditure. We have earlier described that the high-fat diet (HFD) feeding induces an obesity phenotype with high leptin production and alteration of proteins related to endosome sorting, and ubiquitination in the ARC of mice. In this study, real-time PCR data analysis revealed that HFD feeding decreases significantly *Rab3a, Rab3gap1*, and *Rab3gap2* transcript levels in the ARC when compared to the group receiving a control diet. The decrease of *Rab3gap1/2* transcript levels in the ARC was strongly associated with an increase in plasma leptin. Altogether, our studies demonstrate that HFD feeding could be altering the general network of endosome compartmentalization in the ARC of mice, contributing to a failure in exocytosis and receptor recycling.

**Graphical abstract:** 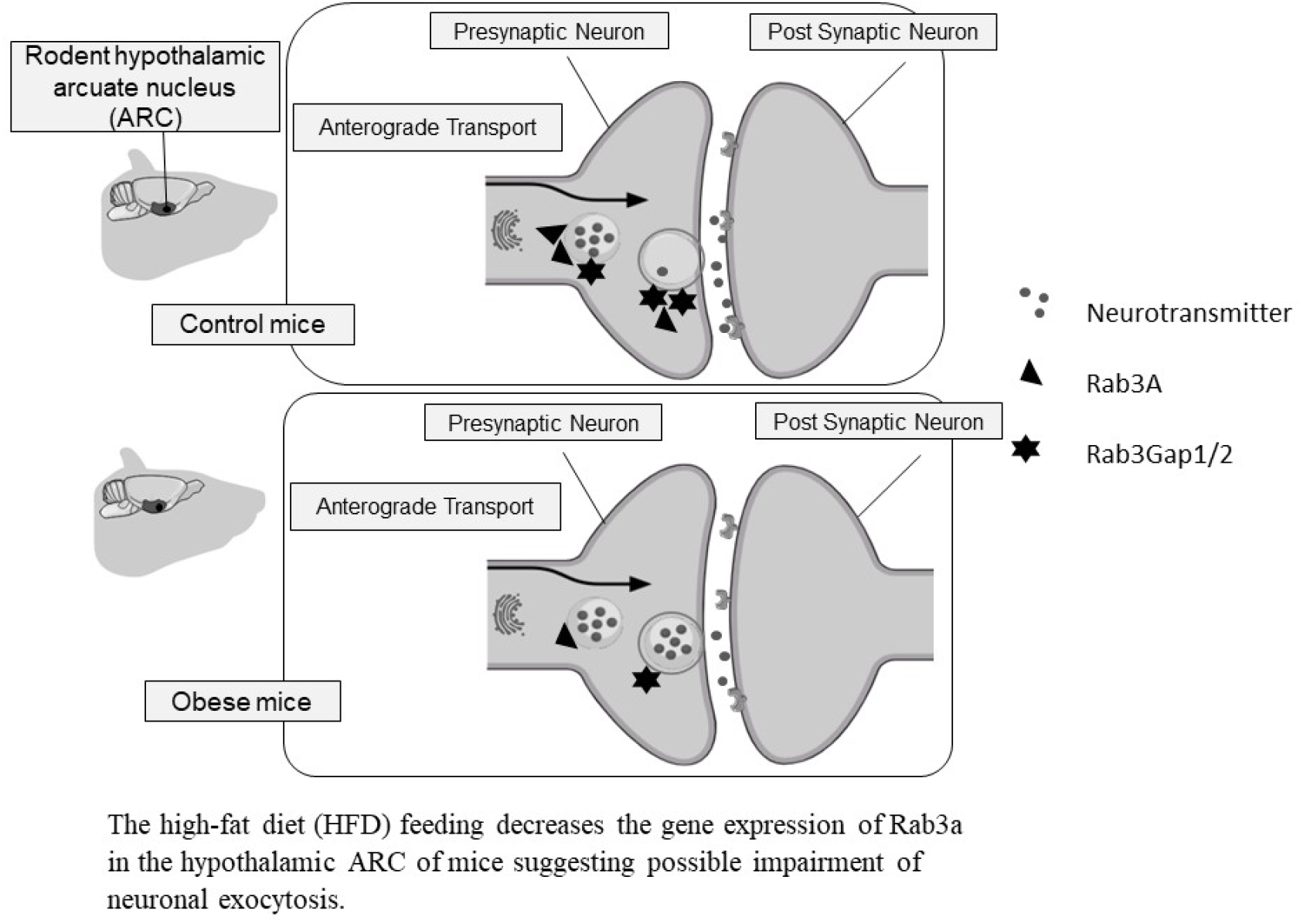

## Introduction

Rabs are small GTP-binding proteins that are generally involved in regulating membrane traffic (Takai et al., 2001). The Rab3A small GTP binding protein is the major Rab protein in the brain and it is described as a regulator of the docking process of vesicles during Ca^+2^-dependent neuronal exocytosis (Geppert et al., 1994a; Oishi et al., 1998; Sedej et al., 2013). Rab3A cycling between GTP-bound and GDP-bound is regulated by the catalytic subunit p130 (Rab3gap1) and the noncatalytic subunit p150 (Rab3gap2) (Fukui et al., 1997; Nagano et al., 1998). Mutations of p130 were related to the pathogenesis of Warburg Micro syndrome (Aligianis et al., 2005). Moreover, it has been described that Rab3gap1 deficiency in mice led to the inhibition of CA1 hippocampal Ca^+2^-dependent glutamate release from cerebrocortical synaptosomes, altering neuronal plasticity (Sakane et al., 2006a). We have previously described that proteins linked to vesicle trafficking are reduced in the ARC of the hypothalamus in HFD mice, as determined by transcript and protein expression of Snx3, Snx27, and Ube2o (Oliveira et al., 2019; Rubatino et al., 2018). In the ARC, the regulation of energy expenditure and food intake is regulated by orexigenic NPY/AgRP (Neuropeptide Y/ Agouti Related Peptide) neurons and by the anorexigenic POMC (Pro-opiomelanocortin) and CART (Cocaine Related Transcripts) neurons (Heisler et al., 2002; Huszar et al., 1997). The exposition of mice to HFD feeding is recognized as inhibiting the satiety performed by the synaptic signaling of POMC/CART neurons in the ARC to another hypothalamic nucleus (Coll et al., 2013). This resultant disbalance of energy expenditure and food intake that leads to obesity is well-recognized to be related to alterations of leptin receptor signaling and repression of POMC expression mediated by low-grade inflammation (Seong et al., 2019). HFD feeding also alters other molecules that control the leptin receptor trafficking and lysosomal degradation in the ARC of mice, contributing to a limited receptor surface exposure and leptin resistance (Roujeau et al., 2019). Other authors recently described that vesicular glutamate and GABA transporters are decreased in the hypothalamus of HFD mice (Lizarbe et al., 2019). All these alterations in the brain of HFD mice are related to the low-grade inflammatory insults and the chronicity of this status. It is known that small GTPases are important for the entire intracellular traffic and synaptic secretion, associating with synaptic vesicular membranes during anterograde transport (Geppert et al., 1994). Given the relevance of Rab3A in the regulation of neurotransmitter and hormone exocytoses, we investigated if HFD feeding alters the transcript levels of *Rab3A, Rab3gap1*, and *Rab3gap2* genes in the ARC of mice.

## Methods

### The murine experimental model of obesity

The high-fat diet obesity model with C57BL/6J^Unib^ was described earlier (Rubatino et al., 2018). All animal procedures were approved by the ethics committee from Ensino e Pesquisa-Santa Casa de Belo Horizonte (EP-SCBH), approval number #0001-15. Four weeks old C57BL/6J^Unib^ were obtained from the Rodents Breeding Center at the Universidade Federal de Minas Gerais (UFMG) and fed a standard chow diet (StD) for 8 weeks. Animals were kept in 30 cm x 18 cm x 13 cm cages (length x width x height) with up to 5 animals per cage in a temperature and light-controlled environment and allowed access to food and water *ad libitum*. Standard diet (StD), 3.7 kcal/g, containing 10% saturated fat, and HFD, 5.3 kcal/g, containing 60% saturated fat, were purchased from Prag Soluções Biociências, Brazil. HFD was introduced when mice were 12 weeks old and maintained for 8 weeks. After this time, mice were euthanized by a trained experimenter using a guillotine to preserve their brains of stress which is accompanied by euthanasia using overdose with xylazine and ketamine. Our HFD obesity murine model presented a similar pattern of weight gain, augment of plasmatic leptin, and hyperglycemia as the previously described diet-induced obesity (DIO) model using C57BL/6J mice (Fontaine and Davis, 2016).

### Determination of transcript levels by real-time PCR in the ARC of the hypothalamus

For the gene expression analysis, the brains were removed shortly after euthanasia by decapitation and immediately immersed in liquid nitrogen. The hypothalamus was dissected with a scalpel and stored in the RNAlater solution. RNA extractions were made using RNeasy Tissue Microarray, (Qiagen®, Hilden, Germany) following the manufacturer’s recommendations. We performed transcript quantifications using a quantitative Real-Time PCR for the genes *Rab3gap1* (NM_178690), *Rab3gap2* (NM_001163754), and *Rab3A* (NM_001166399.2). All samples were run in triplicate. Reactions were performed with ABI 7500 Fast, Applied Biosystems. The fold change values were acquired considering beta-actin (NM_007393) as a housekeeping gene. Data were represented by box plots with median and whiskers of inferior (IL) and superior limit (SL); the outliers (OTL) were calculated using 1^st^ and 3^rd^ quartiles and IQI (interquartile interval) by box plot visualization using R version 4.1.0 software. The Shapiro Wilk normality test was used to determine the distribution of the variables. The *p*-value was determined by Mann-Whitney non-parametric test. The simple linear regression was performed using R version 4.1.0 software. Variable residuals were evaluated by the Shapiro Wilk test to the normal distribution analysis. In the linear regression models, only the weight and Gap1 fold change variables presented non-parametric distribution. The Breusch-Pagan test was used for homoscedasticity, and all variables in the regression models presented homoscedasticity. The presence of outliers was evaluated by graphical representation of the residuals in the linear regression models, and also, by the Durbin-Watson test. No outliers were detected. Summary analysis of the linear regression models provided F-statistic and p – values (plotted in the panels).

## Results and Discussion

### HFD feeding altered Rab3A, Rab3gap1, and Rab3gap2 transcript levels in the ARC of the hypothalamus of mice

This study demonstrates that HFD intake is associated with the reduction of the expression of genes that are essential to the exocytosis of synaptic vesicles in the hypothalamus of mice, which could contribute to energy homeostasis imbalance. The HFD mice presented lower transcript levels of *Rab3a* in the ARC (median, IL, SL; 0.85, 0.03, 0.99 versus 1.02, 0.78, 1.23 fold-change of controls; p=0.003417) (Figure 1a). It is known that the guanine nucleotide forms of Rab3A regulate the transport of vesicles and their docking to the active zone in nerve terminals after Ca^++^-induced depolarization pulse (Geppert et al., 1994b; Leenders et al., 2013; Tsuboi, 2006; van Weering et al., 2007). This activity is determined by the normal cycling between the GDP/GTP forms of Rab3A which are regulated by the GDP dissociation inhibitor (GDI), the GDP/GTP exchange protein (GEP), and the GAPs; also called GTPase activating proteins (Takai et al., 2001). The GTPase activity of GAPs converts the active GTP-form of Rab3A into an inactive GDP-form, suggesting that GAPs are important to the cycling of the Rab3A function in Ca^+2^-triggered exocytosis in the terminal nerve (Sakane et al., 2006b). Indeed, we also found lower transcript levels of *Rab3gap1* (catalytic subunit p130) and *Rab3gap2* (noncatalytic subunit p150) in the ARC of HFD mice when compared to controls (Figure 1b and 1c). The HFD mice presented significantly lower levels of *Rab3gap1* transcripts (median, IL, SL; 0.57, 0.55, 0.59 versus 1.06, 1.103, 1.137-fold change of controls; p=0.01293). The HFD outliers (OTL) were 0.572 and 1.139 (IQI = 0.032, Q1 = 0.97, Q3 = 1.37), the StD OTL were 0.652 and 1.106 (IQI = 0.166, Q1 = 0.59, Q3 = 0.59). The outliers of Gap1 are represented in the Figures. The *Rab3gap2* transcript levels of HFD were also decreased when compared to controls (median, IL, SL; 0.635, 0.623, 0.648 versus 1.246, 0.627, 1.264-fold change of controls; p=0.01908).

**Figure 1:**
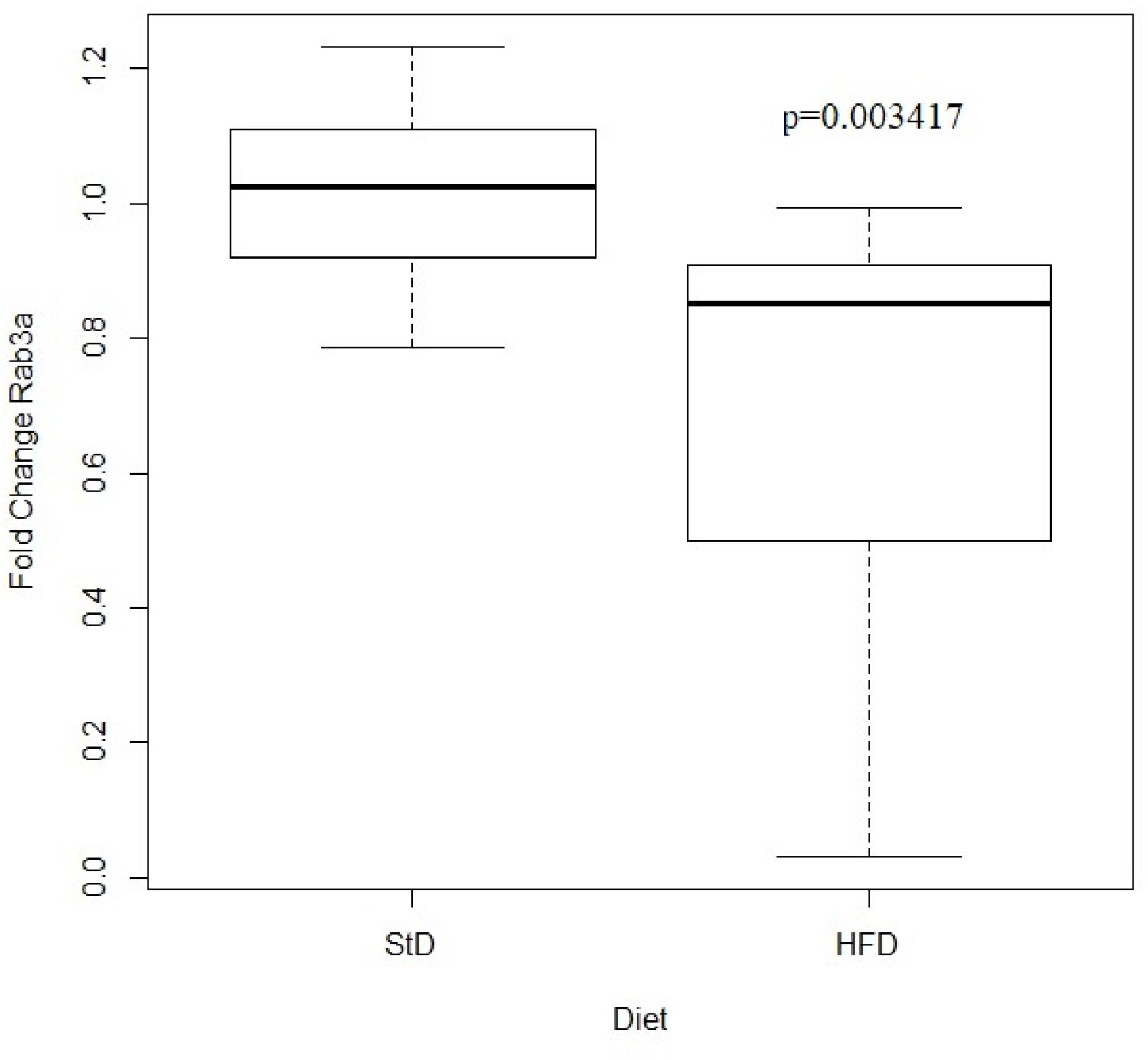
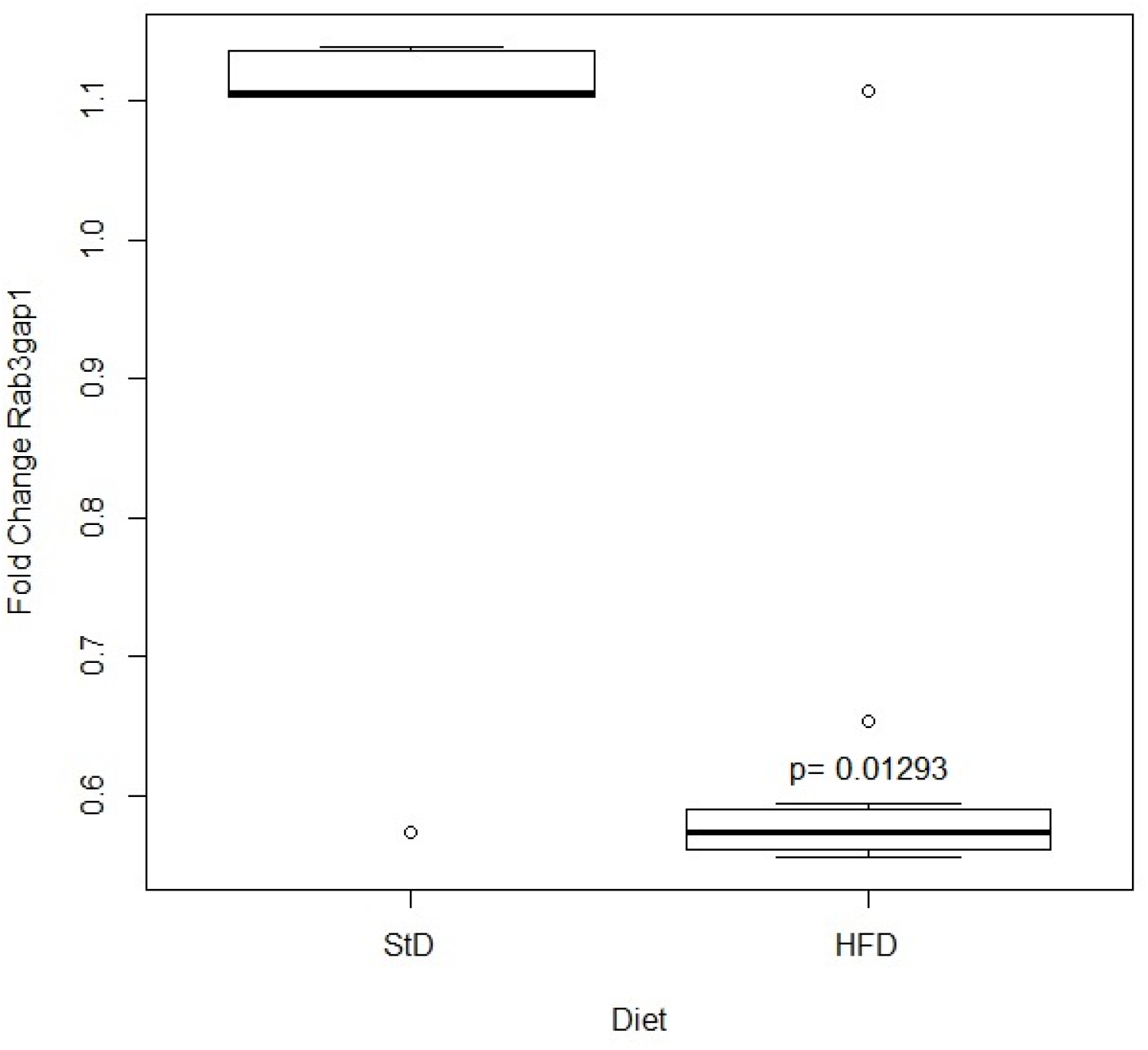
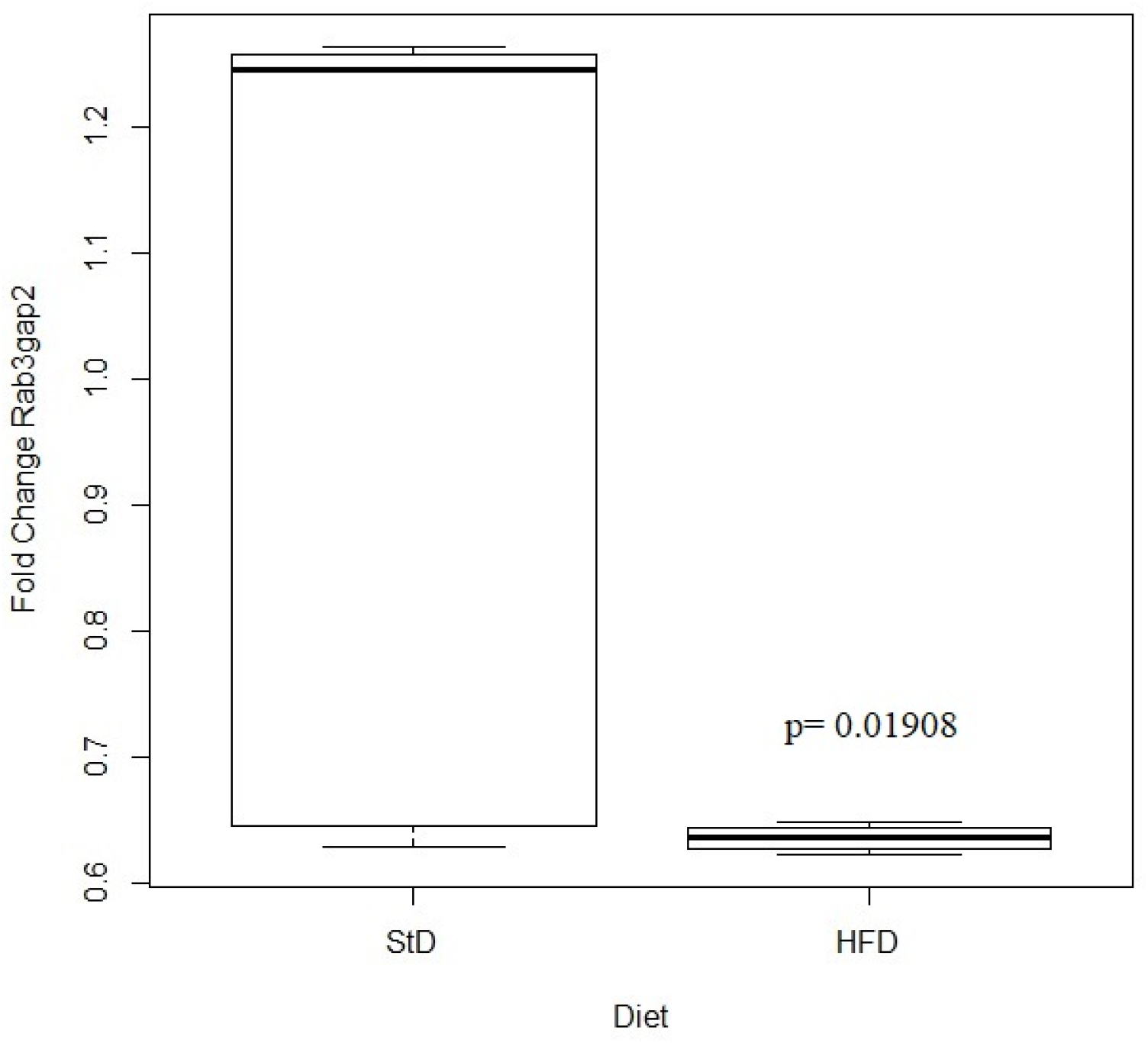
High-fat diet (HFD) decreases Rab3a, Rab3Gap1, Rab3Gap2 transcript levels in the ARC of mice. Twelve-week-old C57BL/6J^unib^ male mice were randomly assigned to a standard (St) (n=6) or HFD (n=6). Gene expression of *Rab3a* (a), *Rab3Gap1*(b), and *Rab3Gap2* (c) data were determined by qPCR on the 8th week and represented as fold change relative quantification. Box plots with whiskers are represented by inferior (IL) and superior limits (SL); outliers (OTL) are represented by open labels in the Gap1 panel. Shapiro-Wilk (W) normality test was used to verify the distribution of variables. Comparisons between groups were evaluated by the Mann-Whitney test, p<0.05 was considered significant. For both groups, n=6 in duplicates. The assay was a representation of three different ones. Statistical analysis and plots were generated by R Version 4.1.0.

Therefore, the inhibition of *Rab3gaps* gene expression observed in the ARC of HFD mice may result in a reduction of GTP hydrolysis, dysregulating the Ca^++^ -dependent exocytosis through Rab3A GTP-bound form accumulation, as described for p130-deficient mice (Sakane et al., 2006c). The reduction of *Rab3gaps* transcript levels was strongly correlated with the augmentation of plasma leptin, glycemia, and total body weight gain (Figure 2). The cause-effect mechanism between diet and gene expressions was not evaluated. However, it is relevant to investigate the possibility that the exocytosis of the neuropeptides α-MSH and CART, as part of the hypothalamic satiety mechanism, may be impaired in the ARC of HFD mice, as previously described (4,20), or whether the relationships among the fattening, satiety, neuropeptide exocytosis, and Rab3 GTPases contribute to obesity.

**Figure 2:**
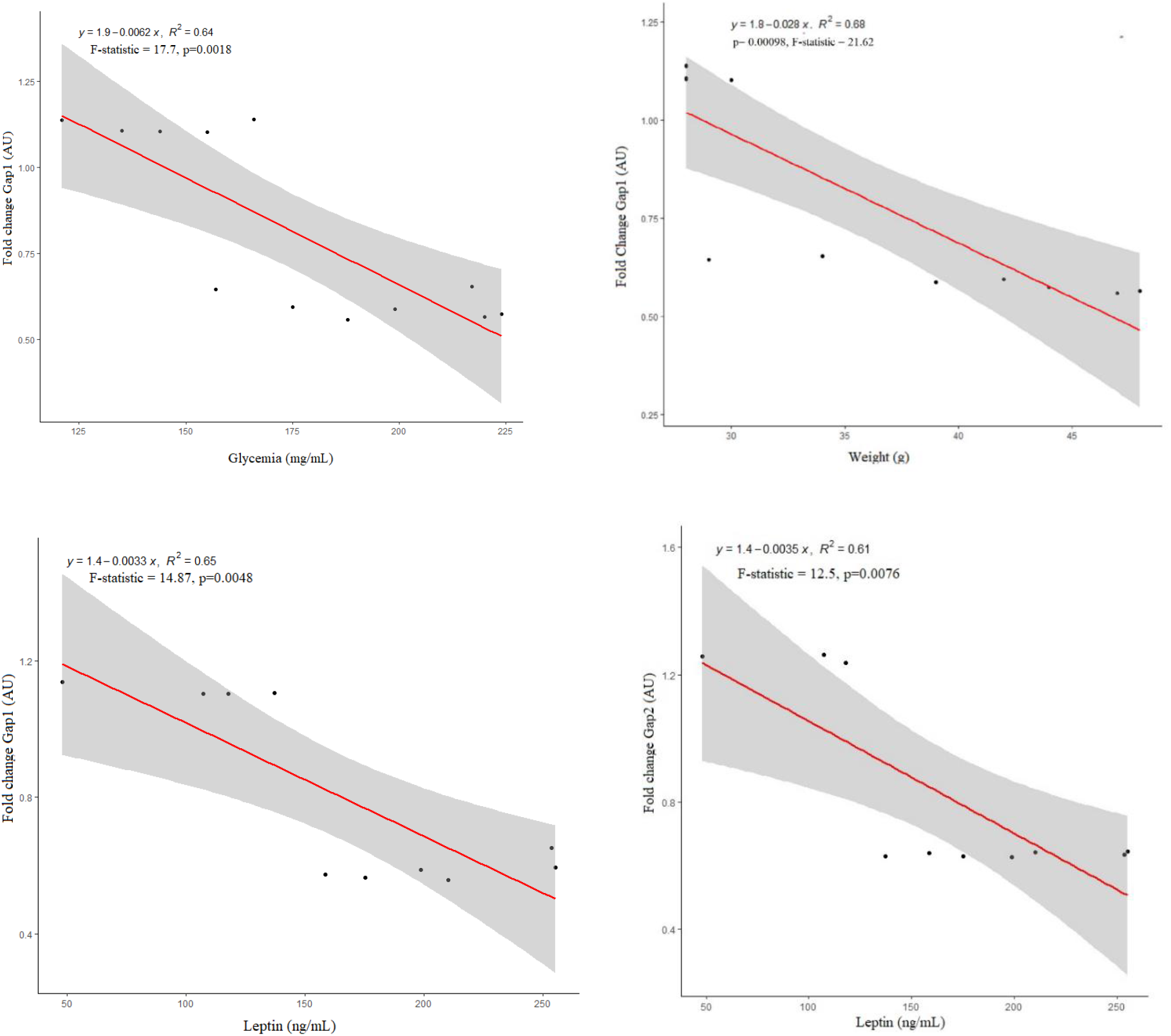
Rab3Gap1 transcript levels in the ARC of HFD mice are negatively associated with fattening, glycemia, and leptin production. Twelve-week-old C57BL/6J^unib^ male mice were randomly assigned to a standard (St) (n=6) or HFD (n=6). In the 8th week, in the absence of fasting, each animal had the body mass and glycemia measured, the gene expression evaluated by qPCR, and plasma leptin determined by ELISA. Gene expression data were represented as fold change relative quantitation (AU). The transcript levels were associated with the total body weight, glycemia, and plasma leptin by simple regression analysis and plotted in panels. The linear regression analysis was performed in R software 4.1.1. Parameters of the models are described in methods. Normal distribution was evaluated by the Shapiro Wilk test, the homoscedasticity by Breusch-Pagan test, and the equation and r^2^ index were plotted in the panels. The F-statistic and the p-value indicate the validity of the linear regression model. Each dot represents one animal. Data are plotted as regression linear lines and 95% confidence interval. p<0.05 was considered significant to the model. The assay was representative of three different ones.

On the other hand, our group has been investigating if the obesity phenotype of mice is associated with alterations in the endosome sorting proteins in the ARC of the hypothalamus, searching for therapeutic targets to regulate satiety (Oliveira et al., 2019; Rubatino et al., 2018). As previously described, we found that in mice receiving HFD there is a reduction of the expression of the sorting nexin type (Snx)3, and the Snx27, which are essential to the retrograde and the fast pathway of recycling vesicles in the cytoplasm, respectively (Oliveira et al., 2019). Besides, we also found that HFD feeding reduces the expression of Ube2O, which is responsible for the ubiquitination of the retromer-wash complex, which leads to endosomal actin nucleation in the cell (Oliveira et al., 2019). It has been described that prolonged exposition of swiss mice to HFD induces ubiquitinated protein accumulation and impairs the process of autophagy in the ARC of these animals (Ignacio-Souza et al., 2014).

Our previous data have also shown that *Tbc1d5* transcript levels were significantly decreased in the ARC of HFD mice (Rubatino et al., 2018). The Tbc1d5 presents GAP activity toward Rab7 inactivation in the retromer complex-mediated endocytosis (Seaman et al., 2009), besides being essential to autophagosome formation (Popovic and Dikic, 2014).

## Conclusion

Together, the findings obtained so far allow us to conclude that the high-fat diet feeding is associated with a reduction of expression of factors involved in the vesicle trafficking pathways and in the exocytosis in the ARC, which contributes to the theory of the disrupted hypothalamic function in obesity.

## Abbreviations

(α-MSH): Alpha-melanocyte-stimulating hormone
(ARC): Arcuate nucleus of the hypothalamus
(CART): Cocaine Related Transcripts
(DIO): Diet-induced obese
(GAP): GTPase activating protein
(GTP): Guanosine-5’-triphosphate
(GDP): Guanosine-5’-diphosphate
(HFD): High-fat diet
IEP- SCBH: Instituto de Ensino e Pesquisa – Santa Casa de Belo Horizonte
(NPY/AgRP): Neuropeptide Y/Agouti Related Peptide
(Snx): Nexins
(POMC): Pro-opiomelanocortin
(RT-PCR): Reverse transcriptase-polymerase chain reaction
(StD): Standard chow diet

## Declarations

### Ethics approval and consent to participate

The present project was submitted and approved by the Animal Research and Experimentation Ethics Committee of Santa Casa de Belo Horizonte, followed by the number #0001-15.

### Consent for publication

All authors had access to the manuscript and approved it for publication.

### Data Availability Statement

The data that support the findings of this study are available from the corresponding author upon reasonable request. We confirm that all results are original and have not been accepted elsewhere.

### Competing interests

The authors declare that they have no competing interests.

### Funding

The study was supported by FAPEMIG grant number APQ-01996-14.

### Author Contributions

All authors contributed to the preparation of the manuscript. LAF, FVMR, KSF, FSJ CJCJ, and AAB accomplished experimental design and literature analysis. LAF, FVMR, MLF, and KSF performed experimental assays and data analysis. KSF, LAF, LRO, and AAB wrote the manuscript. KSF, LAF, FVMR, LRO, and AAB revised the manuscript.

## Acknowledgments

Not applicable

## Notes

### Competing Interest Statement

The authors have declared no competing interest.

